# Seven-year-olds recall non-adjacent dependencies after overnight retention

**DOI:** 10.1101/670257

**Authors:** Gesa Schaadt, Mariella Paul, R. Muralikrishnan, Claudia Männel, Angela D. Friederici

**Affiliations:** Department of Neuropsychology, Max Planck Institute for Human Cognitive and Brain Sciences, Stephanstr. 1a, 04103 Leipzig, Germany; Department of Neurology, Max Planck Institute for Human Cognitive and Brain Sciences, Stephanstr. 1a, 04103 Leipzig and Clinic for Cognitive Neurology, Medical Faculty, Leipzig University, Liebigstr. 16, 04103 Leipzig, Germany; Berlin School of Mind and Brain, Humboldt-Universität zu Berlin, Luisenstraße 56, 10117 Berlin, Germany; Max Planck Institute for Empirical Aesthetics, Grüneburgweg 14, 60322 Frankfurt am Main, Germany

**Author notes:** Correspondence concerning this article should be addressed to Gesa Schaadt, Department of Neuropsychology, Max Planck Institute for Human Cognitive and Brain Sciences, Stephanstr. 1a, 04103 Leipzig, Germany. These authors contributed equally (Gesa Schaadt, Mariella Paul).

**Keywords:** non-adjacent dependencies, ERPs, recall, children, development

## Abstract

Becoming a successful speaker depends on acquiring and learning grammatical dependencies between neighboring and non-neighboring linguistic elements (non-adjacent dependencies; NADs). Previous studies have demonstrated children’s and adults’ ability to distinguish NADs from NAD violations right after familiarization. However, demonstrating NAD recall after retention is crucial to demonstrate a lasting effect of NAD learning. We tested 7-year-olds’ NAD learning in a natural, non-native language on one day and NAD recall on the next day by means of event-related potentials (ERPs). Our results revealed ERPs with a more positive amplitude to NAD violations than correct NADs after familiarization on day one, but ERPs with a more negative amplitude to NAD violations on day two. This change from more positive to more negative ERPs to NAD violations possibly indicates that children’s representations of NADs changed during an overnight retention period, potentially associated with children’s NAD learning. Indeed, our descriptive analyses showed that both ERP patterns (i.e., day one: positive, day two: negative) were related to stronger behavioral improvement (i.e., more correct answers on day two compared to day one) in a grammaticality judgment task from day one to day two. We suggest these findings to indicate that children successfully built associative representations of NADs on day one and then strengthened these associations during overnight retention, revealing NAD recall on day two. The present results suggest that 7-year-olds readily track NADs in a natural, non-native language and are able to recall NADs after a retention period involving sleep, providing evidence of a lasting effect of NAD learning.

**Highlights:**

- 7-year-olds’ non-adjacent dependency learning in a foreign language tested
- Children gave grammaticality judgments while electroencephalography was recorded
- Brain responses revealed children’s learning of non-adjacent dependencies
- Brain responses after overnight retention showed different polarity
- Children’s recall of dependencies after sleep associated with representation change

## 1. Introduction

Language is made up of different building blocks, combined together to form sentences. The grammar of a given language defines the rules for these combinations. For example, grammatical rules define that determiners can be combined with nouns (*The girl*), but not with verbs (\**The give*). Grammatical dependencies can be formed not only between neighboring elements, but also between non-neighboring elements of a sentence. For example, in the sentence *The girlSg smilesg, girlSg* and *-sSg* form a grammatical dependency (i.e., number agreement) that spans one element (smile). In theory, these dependencies can span an arbitrary number of elements, as demonstrated in the following example: *The girlSg who visited us yesterday smilesSg*. These types of dependencies, called non-adjacent dependencies (NADs), are important grammatical rules of a language, such that becoming a proficient speaker and listener of languages highly depends on acquiring these rules (see Wilson et al., 2018).

Adults have been shown to be able to process and learn NADs in a number of behavioral studies (e.g. Frost & Monaghan, 2016; Gómez, 2002; Newport & Aslin, 2004; Peña, Bonatti, Nespor, & Mehler, 2002). For example, Gómez (2002) exposed adults to an artificial language containing NADs in the form of three-syllable strings. In this study, the artificial language learning task consisted of NADs that were realized as AXC structures, with A and C being the dependent elements and X being variable elements. After familiarization to these strings, participants were shown a mixture of strings, either containing familiarized NADs or NAD violations; they were asked to indicate whether a given string followed the rules of the familiarized artificial language. The results showed that adults are in principle able to learn NADs (Gómez, 2002). However, adults’ NAD learning has been shown to be somewhat restricted, as several studies demonstrated that adults only successfully learned NADs when phonological cues between dependent elements were provided (Mueller, Bahlmann, & Friederici, 2008 ; Newport and Aslin, 2004; Peña et al., 2002). Taken together, behavioral studies demonstrated that adults are able to learn NADs in an artificial language. Becoming a successful speaker, however, already starts in early infancy; and even infants have been shown to be able to learn NADs (Gómez and Maye, 2005; Lany & Gómez, 2008). For example, Gómez and Maye (2005) exposed infants to AXC grammars using the Head Turn Preference Procedure (Nelson et al., 1995), with which they measured infants’ looking time towards an auditory stream played on either side of the infant. Specifically, they first familiarized infants with the AXC grammar (i.e., NADs), which was then followed by the presentation of correct or incorrect (i.e., containing a violation) NADs. Fifteen-month-old infants oriented more towards the familiarized stimuli (i.e., correct NADs) than to violations (i.e., incorrect NADs), indicating that 15-month-olds learned the AXC grammar (Gómez & Maye, 2005). However, infants’ NAD learning underlies some restrictions depending on how the NADs are presented (e.g. Höhle, Schmitz, Santelmann, & Weissenborn, 2006; Santelmann & Jusczyk, 1998). For example, 18- to 19-month-old infants can only learn NADs when the intervening elements consist of three syllables or less. If there are more intervening elements, NAD learning breaks down (Höhle et al., Santelmann & Jusczyk, 1998). Taken together, both adults and infants are able to learn NADs in principle. However, the processes underlying NAD learning cannot be fully understood by using offline behavioral methods alone, but should be supplemented by online methods, such as the serial reaction time task or the click detection task (Gómez, Bion, Mehler, 2011; Misyak, Christiansen, Tomblin, 2010). In addition, event-related potentials (ERPs) have been used as an online method to investigate NAD learning. Such online methods allow the more direct examination of the time course of learning and the possible change of underlying learning mechanisms.

Mueller, Oberecker, and Friederici (2009) used ERPs to investigate the learning of NADs that were embedded in natural speech in a foreign language (Italian). During familiarization, they exposed German native speakers, without prior knowledge of Italian, to Italian sentences containing NADs (e.g. “*La sorella sta cantando”*; *the sister is singing*). In testing phases, participants heard a mixture of correct sentences and incorrect sentences containing NAD violations (e.g. “*La sorella sta cantare”*; *the sister is sing*∅Φ). By comparing ERPs to incorrect sentences with ERPs to correct sentences during testing phases, a series of studies could show that both infants (under passive listening conditions, i.e. without a task; Friederici, Mueller, & Oberecker, 2011) and adults (under active conditions, i.e. with a task; Mueller et al., 2009) are able to learn these NADs embedded in a miniature version of Italian. Infants showed a more positive ERP response to incorrect compared to correct NADs, while adults showed a more negative ERP response. Interestingly, when adults’ prefrontal cortex (PFC) was inhibited using transcranial direct current stimulation (tDCS), adults’ ERP response to incorrect compared to correct NADs changed from a negative ERP to a late positive ERP, which was interpreted to indicate different underlying processes (Friederici, Mueller, Sehm, & Ragert, 2013). The late positive ERP found in adults whose PFC was inhibited was similar to infants’ positive ERP responses to NAD violations (Friederici et al., 2011), whose PFC is not yet fully developed (Huttenlocher, 1990). Thus, the polarity difference of the ERP responses to NAD violations seems to be not only due maturational changes between infancy and adulthood, but moreover due to an underlying difference in learning mechanisms. Similarly, studies of language development in early childhood have demonstrated that the polarity of an ERP effect and a developmental change of the ERP effect polarity can be meaningful in terms of later behavior and indicative of different underlying processes (Kooijman, Junge, Johnson, Hagoort, & Cutler, 2013; Schaadt et al., 2015; see also Eimer, Forster, & van Velzen, 2003, and Penney, Mecklinger, & Nessler, 2001, for evidence of a reversal of polarity that is indicative of behavior in adults).

As indicated by a difference in ERP polarity of components elicited by NAD violations, infants and adults might use different learning mechanisms and develop different representations of the NADs. Specifically, it has been suggested that infants learn NADs more automatically than adults do, also reflected in infants’ ability to learn under passive listening, which adults struggle to do (Mueller, Friederici, & Männel, 2012). Interestingly, Mueller, Friederici, and Männel (2018) showed that children up to the age of 2 years are able to learn NADs under passive listening conditions, while 4-year-olds, similar to adults (Mueller et al., 2012) struggle to do so and may need active task conditions. It has been suggested that the specific need for an active task is associated with a switch in learning mechanisms from associative, bottom-up learning (allowing learning under passive listening conditions) to controlled, top-down learning (hindering learning under passive listening conditions, but facilitating learning under active task conditions; see Skeide & Friederici, 2016). This switch may be associated with PFC maturation (Skeide & Friederici, 2016), which reaches near adult-like maturity around the age of 7 years (Huttenlocher, 1990). While this claim has not been tested longitudinally, there is some evidence for this from the tDCS (Friederici et al., 2013) and the cross-sectional study (Mueller et al., 2018) described above. Taken together, NAD learning mechanisms change during development, which may possibly be linked to PFC development.

Although previous studies (Friederici et al., 2011; Mueller et al., 2009; 2012) convincingly demonstrated that individuals can differentiate familiarized NADs from NAD violations, NAD learning was always tested on the day of the familiarization itself, either on the same items as during familiarization (e.g., Friederici et a., 2011; Mueller et al., 2009) or on novel items (i.e. items that share the same structure as familiarized items, but use different tokens; e.g., Gómez, Bootzin, & Nadel, 2006). This testing procedure provides a measure of whether participants have formed a representation of the familiarized items, which can then be compared to test items. Test items perceived as similar to familiarization items would then be interpreted as adhering to the (possibly unknown) rule. On the other hand, test items judged as dissimilar would be interpreted as not adhering to the underlying rule (similarity-based learning; see Opitz & Hofmann, 2015). While testing NAD learning on the same day of familiarization is certainly informative, it is a matter of discussion whether this should be interpreted as evidence that the underlying rules have been learned, rather than some surface-based features of the NADs. This is because the knowledge of the underlying rules that characterize the (artificial) grammar only builds up over time (Opitz & Hoffman, 2015) and might not be fully present immediately after a relatively brief familiarization. Thus, it is important to retest NAD learning after a period of time in order to investigate whether NAD learning had a lasting effect and to show that learned NADs are not simply forgotten again shortly after familiarization. In order to investigate whether participants have really learned the underlying rules and could recall them after a period of time, several studies have investigated recall of grammatical rules after a retention period. For example, Fischer, Drosopoulos, Tsen, and Born (2006) investigated the effect of a retention period on artificial grammar learning in adults. The authors showed that before sleep there was no evidence for above-chance level performance in adults in a generation task, during which participants had to predict the next letter in a string based on the artificial grammar. However, after a retention period involving sleep participants could solve the task successfully, which was not the case after a retention period without sleep. A number of studies has demonstrated that this benefit of a retention period involving sleep is linked to a change in representations (see Diekelmann, & Born, 2010; Ellenbogen, Hu, Payne, Titone, & Walker, 2007; Fischer et al., 2006; Wagner, Gais, Haider, Verleger, & Born, 2004). Davis and Gaskell (2009) suggested that a model of memory consolidation, the complementary learning systems model, could also apply to the linguistic domain, specifically word learning. Under this model, new knowledge, such as a newly encountered word, is initially stored in episodic memory, where it is not yet integrated into the lexicon. New words are then consolidated into lexical memory over time, facilitated by sleep (Henderson et al., 2012; Smith et al., 2016; Tamminen et al., 2010). Especially infants and children were shown to benefit from a retention period (particularly when retention involved sleep; Backhaus, Hoeckesfeld, Born, Hohagen, & Junghanns, 2008; Henderson et al., 2012; Hupbach, Gómez, Bootzin, & Nadel, 2009; Smith et al., 2016) and for generalizing learned information to new input (Gómez, Bootzin, & Nadel, 2006). A study by Friedrich, Wilhelm, Mölle, Born, and Friederici (2017) linked a change in representations of learned associations during the course of a retention period to particular ERPs. In this study, infants were exposed to object-word pairs followed by a retention period that either involved a long nap, a short nap, or no sleep. Before retention, there was no evidence for learning of the object-word pairs and neither did the group without sleep show any sign of learning after retention. In contrast, infants who had a short retention period (30 minutes on average) involving sleep showed consolidation of the object-word pairs. However, the ERPs only revealed a late negativity, which was interpreted to be indicative for a phonological association between the word and object, but not for a lexical-semantic representation of the object-word pairs in long-term memory. Only those children who had a longer consolidation period (50 minutes on average) involving sleep also showed ERP evidence of lexical-semantic representations of word meaning in long-term memory in form of an N400 (i.e., earlier negativity; Friedrich et al., 2017). Thus, this study demonstrates that children benefit from a retention period involving sleep, which most likely leads to the ERP effects of successful recall of learned associations after the retention period. Given these promising findings showing a beneficial effect of a retention period involving sleep on long-term memory consolidation, we aimed at investigating the effect of retention involving sleep on the recall of NADs as important grammatical rules of language.

Thus, in the present ERP study, we investigated 7-year-old children’s recall of NADs embedded in a miniature version of a foreign language (i.e., Italian), using the same paradigm as Mueller and colleagues (2009), including a grammaticality judgment task. We invited our participants on two consecutive days, ensuring a retention period involving sleep to test recall of NADs. If we can show recall of NADs on day two, we provide evidence that children learned the NADs and that this learning had lasting effects beyond the familiarization day, which goes over and above showing processing differences between correct and incorrect NADs on the same day when familiarization took place. We tested 7-year-olds because they have been shown to be able to successfully perform offline behavioral tasks assessing statistical learning (Raviv & Arnon, 2018; Shufaniya & Arnon, 2018), most likely associated with 7-year-olds’ advanced PFC maturation (Huttenlocher, 1990), playing a crucial role in NAD learning (Friederici, Mueller, Sehm, and Ragert, 2013).

A number of recent studies have raised concerns that group-level offline tasks, which assess statistical learning, may not provide reliable measures of individual differences (Siegelman, Bogaerts, & Frost, 2017; West, Vadillo, Shanks, & Hulme, 2018), particularly in children (Arnon, 2019). Siegelman, Bogaerts, Christiansen, and Frost (2017) suggest that online measures may circumvent some of the problems seen in the reliability of offline tasks. Here, we use ERPs as an online test of NAD learning both at the group level and the individual level, as ERPs have been shown to be a reliable measure of interindividual differences in a variety of paradigms (Cassidy, Robertson, & O’Connell, 2012).

According to the procedure of Mueller and colleagues (2009) in adults, children listened to only correct stimuli (i.e., Italian sentences) during the four learning phases on the first testing day. Each learning phase was followed by a testing phase, during which children listened to incorrect stimuli containing NAD violations intermixed with correct stimuli following the familiarized NAD rule. During the testing phases, children were required to behaviorally indicate whether or not a given stimulus belonged to the language they were familiarized with in the learning phases (i.e., grammaticality judgment task). On the following day, we tested recall of NADs by asking children to perform only the four testing phases, again including the grammaticality judgment task. To capture consolidation and recall of NADs on the next day, we specifically focused on the change in behavior from day one to day two. Successful recall of NADs will be reflected in behavioral improvement from day one to day two (i.e., more correct grammaticality judgments on day two compared to day one). If children learn the NADs on day one and recall them on day two, we expect that children’s ERP responses on both days are associated with their improvement in the number of correct grammaticality judgments from day one to day two. While we will treat this correlational analysis as an exploratory analysis due to reliability concerns (see Siegelman et al., 2017a), linking ERPs to the behavioral outcome may strengthen the interpretability of our results.

## 2. Materials and Methods

### 2.1 Participants

For the present experiment, 49 children were invited. The datasets of 36 children (20 boys) with a mean age of 7.22 years [*Standard Deviation (SD)* = 0.36] entered the final analyses (i.e., the datasets of 13 children were excluded due to movement and perspiration artifacts in the EEG). Children visited the first and second school grade. All participants were German monolinguals and none of the children had any known hearing deficits or neurological problems. In order to ensure that the Italian sentences used for the present study were foreign to the children and thus, functioned as an “artificial” language, we asked the parents about the child’s experience with foreign languages and specifically with the Italian language. One of the 36 children visited a bilingual French-German kindergarten and at school, 11 of the 36 children learned a second language, with two children learning French and nine children learning English. Thus, none of the children had any specific experience with the Italian or Spanish (Spanish and Italian consist of the same NADs) language.

The study followed American Psychological Association (APA) standards in accordance with the declaration of Helsinki from 1964 (World Medical Association, 2013) and was approved by the ethics committee of the University Leipzig. Parental written consent was obtained after children and parents had been informed about the procedure and agreed to participation.

### 2.2 Stimulus material

Mueller and colleagues (2009) provided the stimuli for the present study. They consisted of simple Italian sentences, containing an NAD between an auxiliary and a main verb’s suffix. Sentences were made up of one of two noun phrases (*il fratello*, the brother; *la sorella,* the sister), one of two auxiliaries (*può*, to be able to, first person singular; *sta*, to be, first person singular), and one of 32 verbs. Verbs could either occur in infinitive (e.g., *arrivare*) or in gerund form (e.g., *arrivando*). Between the auxiliary and the verb suffix was a non-adjacent grammatical dependency, such that the auxiliary *sta* required the gerund form *-ando* and the auxiliary *può* required the infinitive form *-are*. In total, 128 correct sentences were generated. All correct sentences were spoken by a female native Italian speaker and digitally recorded. Subsequently, the auditory material was segmented and normalized using the ReZound software. Incorrect sentences were produced by combining auxiliaries with the incorrectly suffixed verbs from a different, correct sentence. This was done by a cross-splicing procedure at the beginning of each verb. In each sentence, the verb was thus exchanged with a verb from a different sentence. To control for splicing effects across conditions, correct sentences were spliced in the same manner.

### 2.3 Experimental Procedure

Participants were invited for two consecutive days. On the first testing day, participating children and their parents were verbally informed about the procedure. Children were asked to provide consent to participate and parents gave written informed consent on behalf of their children. Participating children were read a cover story about an explorer hearing sentences in a foreign language and who needs help deciding whether the words in the sentences fit together. Children were not explicitly told that they were supposed to learn an underlying rule, but to carefully listen to the sentences and decide whether the words in the sentence fit together (for further details, see the exact instructions at https://osf.io/b3e5a/). Further, children were informed that they would be re-invited for the next day, but not that they would be tested on the same grammar again. Our experiment on the first day comprised four alternating learning and testing phases. In each learning phase, participants were presented with 64 correct sentences (256 in total across all learning phases). After a learning phase, a testing phase followed where participants were presented with correct sentences and incorrect versions of the sentences containing NAD violations. Each testing phase consisted of 8 correct and 8 incorrect sentences (64 sentences across all four testing phases). Please note that each testing phase contained different auxiliary-verb-suffix-triplets compared to the preceding learning phase to ensure that participants learned NADs (and not auxiliary-verb-suffix triplets).

For the ERP experiment, participants sat in a sound-attenuated booth in front of a computer screen and stimuli were presented via loudspeaker using Presentation^®^ software Version 14 (Neurobehavioral Systems Inc., Berkely CA, USA). Children were instructed by using a cover story, where they were asked to support an adventurer, needing help to decide whether the sentences in a foreign language are correct or incorrect. Further, they were told that it is important to listen carefully, because otherwise the adventurer would not be able to continue his journey around the world. After the instruction, the experiment started with the first learning phase, in which participants passively listened to the correct NAD sentences. A fixation cross was continuously presented in the middle of the screen to reduce extreme eye-movements. Sentences were presented in a pseudo-randomized order, such that each sentence beginning (i.e., *il fratello sta, la sorella sta, il fratello può, la sorella può*) was not presented more than three-times in a row; and such that a verb could only be repeated every third sentence. From the beginning of each sentence to the beginning of the following sentence, there was an inter-stimulus-interval (ISI) of 3000 ms. Each learning phase (in total 4 learning phases) was followed by a grammaticality judgment task to test for learning effects, with pseudo-randomized presentation following the above mentioned criteria for pseudo-randomization and pseudo-randomized presentation of incorrect and correct sentences, such that correct or incorrect sentences could only be presented twice in a row. Children were required to give grammaticality judgments on each stimulus (i.e., correct vs. incorrect) by using a button-press response device. The trials started with a fixation cross that was presented for 1000 ms, before one of the correct or incorrect sentences was presented. After an ISI of 3000 ms, the simultaneous display of a happy (indicating correct) and a sad face (indicating incorrect) prompted participants to judge the grammatical correctness of the sentence via the provided response keys. The response key assignment (right / left) to the answer type (correct / incorrect) during the testing phases was counterbalanced across participants. Each learning phase lasted for about 3.5 min and each testing phase lasted for about 10 min (i.e., depending on the child’s response times), summing up to a total experimental time of around 60 min.

For the second testing day, our experiment only comprised four testing phases (i.e., grammaticality judgment task as described above), but no learning phases, to investigate NAD learning after a retention period including sleep. Stimuli used for the second testing day were not identical to those used on the first testing day (i.e., sentence beginnings and verbs forming correct and incorrect sentences were combined differently on day two compared to day one; for a list, see https://osf.io/b3e5a). As participants were not presented with the learning phases on the second testing day, the total experimental time was reduced (i.e., around 40 min). EEG was recorded during the whole experiment on both testing days. Behavioral data (i.e., error rates; response times) were recorded for each participant during testing phases on testing day one and testing day two.

### 2.4 EEG Recordings and Analysis

Continuous EEG was recorded with an EGI (Electrical Geodesics, 1998) 128-electrode array (see Figures 1, 2, and 3 for schematic illustration). The vertex (recording site Cz) was chosen as online reference. For the EGI high input impedance amplifier, impedances were kept below 75 kΩ. The sampling rate was 500 Hz and all channels were pre-processed on-line by means of 0.01 – 200-Hz band-pass filter. In addition, vertical and horizontal eye movements were monitored with a subset of the 128 electrodes.

**Figure 1.**
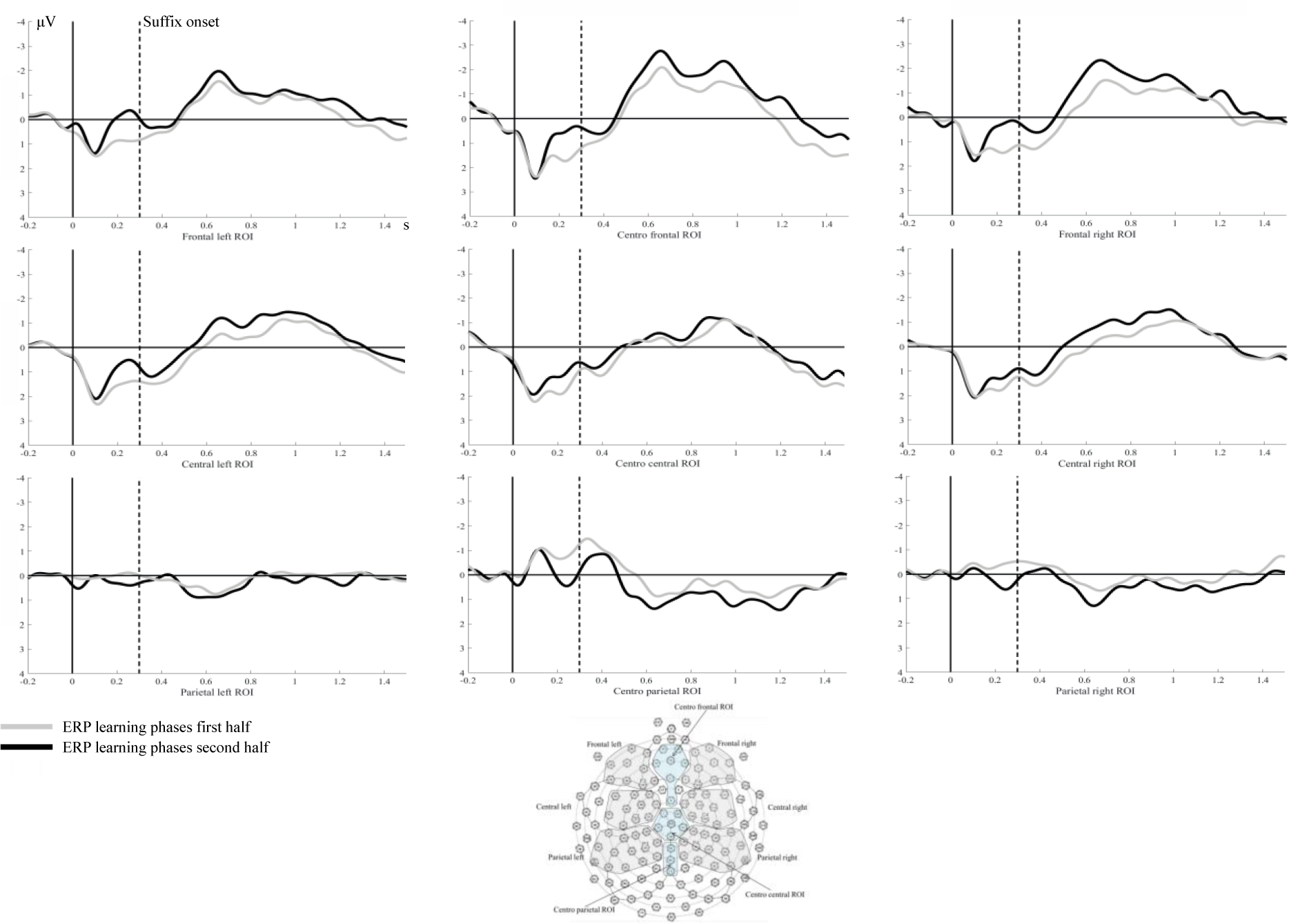
Event-related potentials (ERPs) of the learning phases on day one. Illustrated are the mean ERPs in response to the correct suffixes during the first half of the learning phases (grey line) and to the correct suffixes during the second half of the learning phases (black line) averaged for left, middle, and right frontal, central, and parietal regions of interest (ROIs; see schematic head for details on electrodes).

For off-line EEG analysis, we used the Fieldtrip toolbox for EEG/MEG analysis (Oostenveld, Fries, Maris, & Schoffelen, 2011) and the MATLAB^®^ version R2017b (The MathWorks Inc., 2017). Before preprocessing, EEG data was manually scanned for electrodes with bad or missing signal. Those electrodes were excluded from the respective data set. Note, however that the number of excluded electrodes never exceeded 6 out of 128 (i.e., < 5%) and that excluded electrodes differed across participants. Thereafter, data were off-line re-referenced to the average of all EEG electrodes. Before data were filtered, the sampling rate was reduced to 250 Hz. We then applied a digital low-pass filter of 30 Hz (Kaiser-windowed finite-impulse response low-pass filter, half-amplitude cutoff (-6 dB) of 30 Hz, transition width of 5 Hz) to remove muscle artifacts and a high-pass filter of 0.3 Hz (Kaiser-windowed finite-impulse response high-pass filter, half-amplitude cutoff (-6 dB) of 0.3 Hz, transition width of 0.3 Hz), to remove very slow drifts. In a next step, we extracted trials of –200 to 2000 ms time-locked to the onset of the critical verb (i.e., containing either the correct or incorrect suffix). Across all remaining trials, we identified muscle artifacts with a distribution-based identification approach. We set the rejection threshold to z = 7.0. Trials were visually scanned and, if applicable, further trials with severe artifacts were manually marked and removed. To remove eye-movement artifacts, an independent-component analysis (ICA; Makeig, Bell, Jung, & Sejnowski, 1996) was performed. ICA components were visually scanned and eye movement-related components removed. Before individual averages were computed (baseline corrected from –200 to 0 ms relative to verb-onset), the removed electrodes with bad or missing signal were interpolated by using spherical spline interpolation (Perrin, Pernier, Bertrand, & Echallier, 1989). In a second step, grand averages were computed in relation to the suffix onset for the learning phases (separately for the first and second halves of the experiment) and for the testing phases on day one and day two, separately for verbs containing correct suffixes (i.e., NAD was not violated) and verbs with incorrect suffixes (i.e., NAD was violated).

### 2.5 Statistical Analysis

For statistical analyses, we used the Statistical Package for the Social Sciences (SPSS) Software Version 24 (IBM; Walldorf, Germany).

#### 2.5.1 Behavioral data

For each testing day, statistical means of response times (RTs) in ms and correct answers in percent were calculated for each participant. To analyze whether RTs and correct answers differed between day one and day two, we calculated dependent *t-*Tests. In a next step, we performed binomial tests for each child to determine whether performance (i.e., correct answers in percent) was above chance level in the grammaticality judgment NAD task.^1^.

According to the performed binomial test, the threshold indicating above chance-level performance was 58.2 or more correct answers in percent (*p* < .05), which we used to classify each child’s grammaticality judgment NAD task performance. Finally, we obtained a score indicating whether children’s task performance changed (i.e., number of correctly answered trials) from day one to day two by calculating the difference between correct answers on day two and the correct answers on day one.

#### 2.5.2 EEG data

To statistically analyze the ERP data, we defined two frontal regions of interest (ROIs), two central ROIs, and two parietal ROIs for each hemisphere (i.e., left and right; see Figures 1, 2, 3 and Luu & Ferree, 2005). Further, we defined ROIs for the midline (see Figures 1, 2, 3 and Luu & Ferree, 2005). ERP analyses were performed on six time windows (TW) of 200 ms each. The suffix-onset (*-are* and -*ando*) served as criterion for TW definition, as it is the earliest point at which a correct sentence can be distinguished from an incorrect sentence. On average, suffix onset occurred at 267 ms (range: 138 – 408 ms) relative to the onset of the verb stem, such that we defined the first TW of interest to start 300 ms after verb onset.

To identify significant ERP effects of learning across the experiment on day one, we contrasted ERPs in response to the critical suffixes during the first and second halves of the experiment. In order to do so, we calculated a three-factorial repeated measures analyses of variance (ANOVA) with the within-subject factors learning phase (first half, second half), region (left frontal, centro frontal, right frontal, left central, centro central, right central, left parietal, centro parietal, right parietal), and TW (300–500 ms, 500–700 ms, 700–900 ms, 900– 1100 ms, 1100–1300 ms, 1300–1500 ms). If effects involving the factor learning phase reached significance (*p < .*05), post-hoc pairwise comparisons were computed, *p-*values were Bonferroni-corrected and reported p-values are adjusted for multiple testing.

To identify significant ERP differences between the processing of correct and incorrect suffixes during testing phases and whether these potential ERP differences change from day one to day two, we calculated a four-factorial ANOVA with the within-subject factors testing day (day one, day two), condition (correct, incorrect), region (left frontal, centro frontal, right frontal, left central, centro central, right central, left parietal, centro parietal, right parietal), and TW (300–500 ms, 500–700 ms, 700–900 ms, 900–1100 ms, 1100–1300 ms, 1300–1500 ms). Two hundred-ms time windows were chosen to enable comparability to previous studies using the same stimuli (specifically Friederici et al., 2011, who used 200-ms analysis time windows) ^2^. If effects involving the factor condition reached significance (*p < .*05), post-hoc pairwise comparisons were computed, *p-*values were Bonferroni-corrected and reported p-values are adjusted for multiple testing.

In a further step, we analyzed whether significant ERP effects could predict the change in task performance from day one to day two. First, we calculated the ERP difference waves between those contrasts, for which the above-described ANOVAs revealed statistically significant effects (e.g., ERP to incorrect suffixes – ERP to correct suffixes). By calculating such ERP difference waves, we were able to determine the quantity (i.e., difference in amplitude) and quality (i.e., polarity) of potential processing differences, which might be associated differently with task performance. Both, the amplitude and polarity of the difference wave are indicative of the underlying processes involved in stimulus processing (Luck, 2004) and might thus influence task performance differently. For example, Kooijman and colleagues (2013) showed that infants’ polarity of ERP components elicited during a word segmentation task was predictive of later vocabulary. Second, we then calculated a correlational analysis between the ERP difference waves (i.e., for those contrasts that revealed statistically significant effects on day one and on day two) and the change in task performance from day one to day two (correct answers day two – correct answers day one).

## 3. Results

### 3.1 Behavioral Results

Correct answers in percent on day one (*mean* = 48.65%; *SD* = 5.90) did not differ significantly from correct answers on day two (*mean* = 50.73%; *SD* = 6.18; *t* (35) = -1.49; *p* = .15). RTs were significantly shorter on day two (*mean* = 7489.13 ms; *SD* = 3843.89) compared to day one (*mean* = 11552.69 ms; *SD* = 4847.99; *t* (35) = 5.77; *p* < .001). When using the criterion of the binomial test (i.e., when z was set to 1.64, resulting in a above-chance level threshold of 58.2% correct answers in percent), we could identify three children performing above chance level on day one and three children on day two (only partially overlapping; i.e., one child). Further, the *mean* change in behavior was 2.09% (*SD* = 8.41; min = -15.65; max = 17.15), indicating that some children showed more correct answers on day two compared to day one (i.e., positive values) and some children showed less correct answers on day two compared to day one (i.e., negative values).

### 3.2 EEG Results

#### 3.2.1 Learning phases day one

Neither the main effect of learning phase [*F* (1, 35) = 0.41; *p* = .53], nor any interaction involving the factor learning phase [learning phase * TW: *F* (5, 175) = 0.63; *p* = .59, learning phase * region: *F* (8, 280) = 0.71; *p* = .55, learning phase * TW * region: *F* (40, 1400) = 0.92; *p* = .48] reached significance (see Figure 1). Thus, ERPs in response to the critical suffixes during the first half of the experiment did not differ significantly from the ERPs in response to the critical suffixes during the second half of the experiment.

#### 3.2.2 Testing phases day one and day two

We found a significant interaction between the factors testing day and condition [*F* (1, 35) = 4.19; *p* = .049; η^2^ = .11], which could be explained by a more positive ERP response to incorrect suffixes compared to correct suffixes on day one (*p* = .026). Further, we found a significant interaction between the factors testing day, condition, TW, and region [*F* (40, 1400) = 2.03; *p* = .03; η^2^ = .06], which could be explained by a more positive ERP response to incorrect suffixes compared to correct suffixes on day one for the TW 1100–1300 ms at the centro frontal region (*p* = .02), the left frontal region (*p* = .004), and the left central region (*p* = .04) (see Figure 2); and by a more negative ERP response to incorrect suffixes compared to correct suffixes on day two for the TW 900–1100 ms at the left frontal region (*p* = .05) and at the left central region (*p* = .01); and for the TW 1100–1300 ms at the left frontal region (*p* = .02) (see Figure 3).

**Figure 2.**
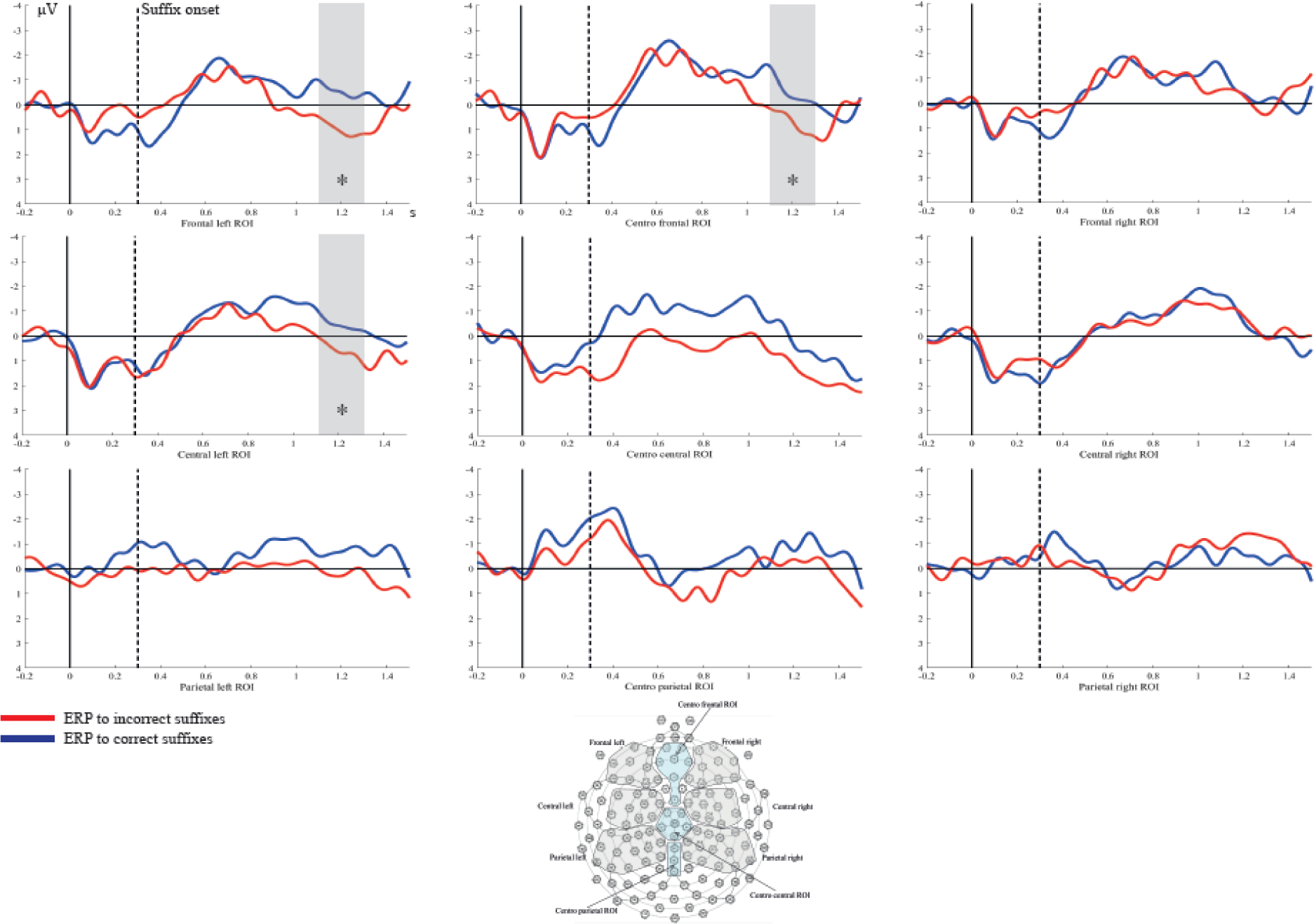
Event-related potentials (ERPs) of the testing phases on day one. Illustrated are the mean ERPs in response to the correct suffixes containing the nonadjacent dependency (blue line) and to the incorrect suffixes violating the nonadjacent dependency rule (red line) averaged for left, middle, and right frontal, central, and parietal regions of interest (ROIs; see schematic head for details on electrodes). Grey bars and asterisk indicate time windows and regions with significant differences between the ERPs of the two conditions (* p < .05).

**Figure 3.**
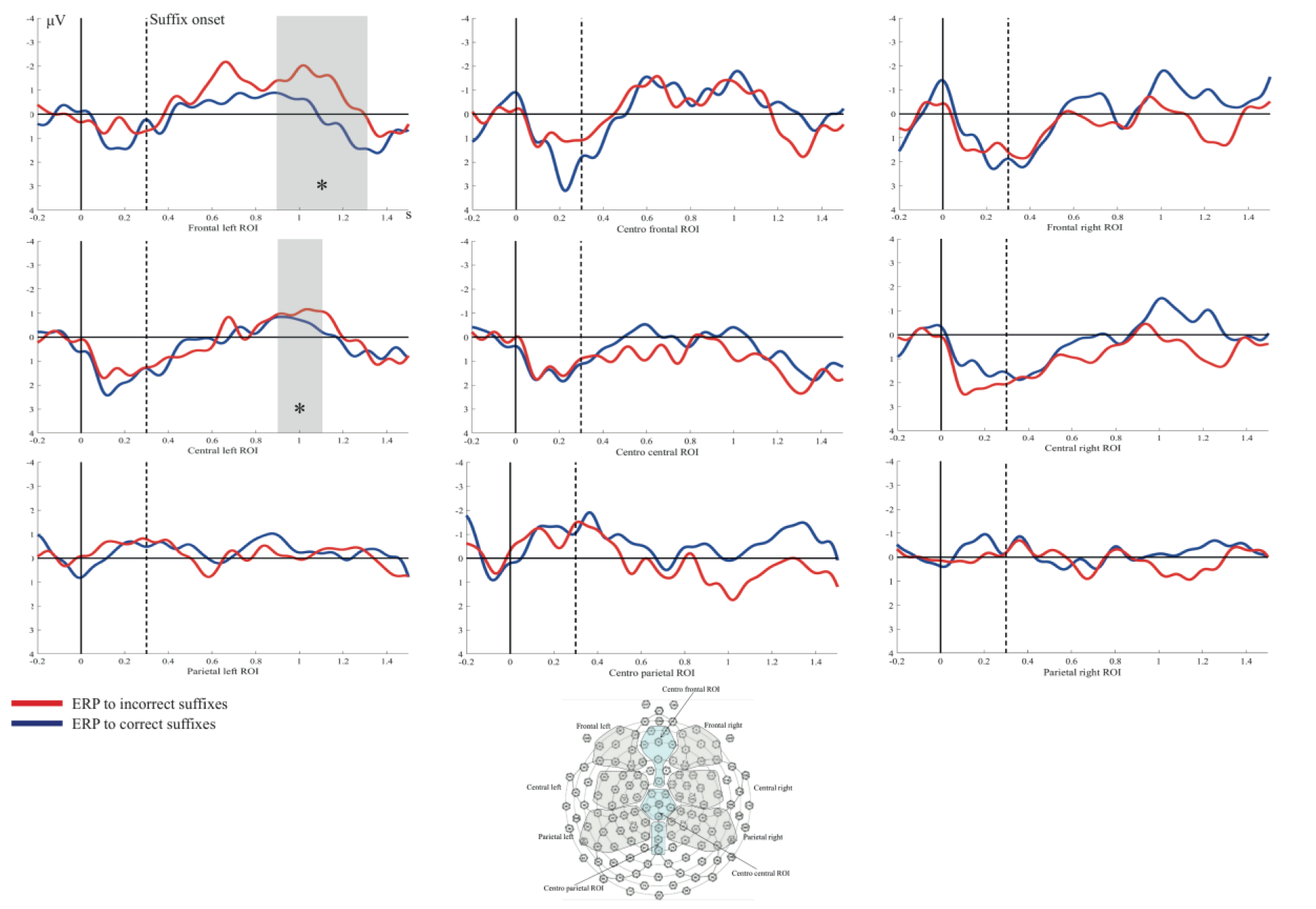
Event-related potentials (ERPs) of the testing phases on day two. Illustrated are the mean ERPs in response to the correct suffixes containing the nonadjacent dependency (blue line) and to the incorrect suffixes violating the nonadjacent dependency rule (red line) averaged for left, middle, and right frontal, central, and parietal regions of interest (ROIs; see schematic head for details on electrodes). Grey bars and asterisk indicate time windows and regions with significant differences between the ERPs of the two conditions (* p < .05).

Thus, we found a more positive ERP response to incorrect compared to correct suffixes between 1100 and 1300 ms, that is, between 800 and 1000 ms after suffix onset on day one and a more negative ERP response to incorrect compared to correct suffixes between 900 and 1300 ms, that is, between 600 and 1000 ms after suffix onset on day two.

### 3.3 Descriptive analyses of behavioral changes in relation to ERPs

Because task performance was not above-chance at the group level and only very few children performed above-chance on an individual level (see section 3.1), we will only analyze the association between behavioral changes and ERPs descriptively, and will refrain from performing inference statistics for this analysis. We calculated Pearson’s bivariate correlation coefficient to analyze the association between ERP difference waves (ERP to incorrect suffixes – ERP to correct suffixes) of those contrasts that revealed statistically significant effects on day one and on day two and the change in task performance from day one to day two (correct answers day two – correct answers day one).

The results showed that the ERP effect between 1100 and 1300 ms on day one (i.e., positivity) was positively associated with the change in task performance from day one to day two (*r* = .21), while the ERP effect between 900 and 1300 ms on day two (i.e., negativity) was negatively associated with the change in task performance from day one to day two (*r* = – .45) (see Figure 4).

**Figure 4.**
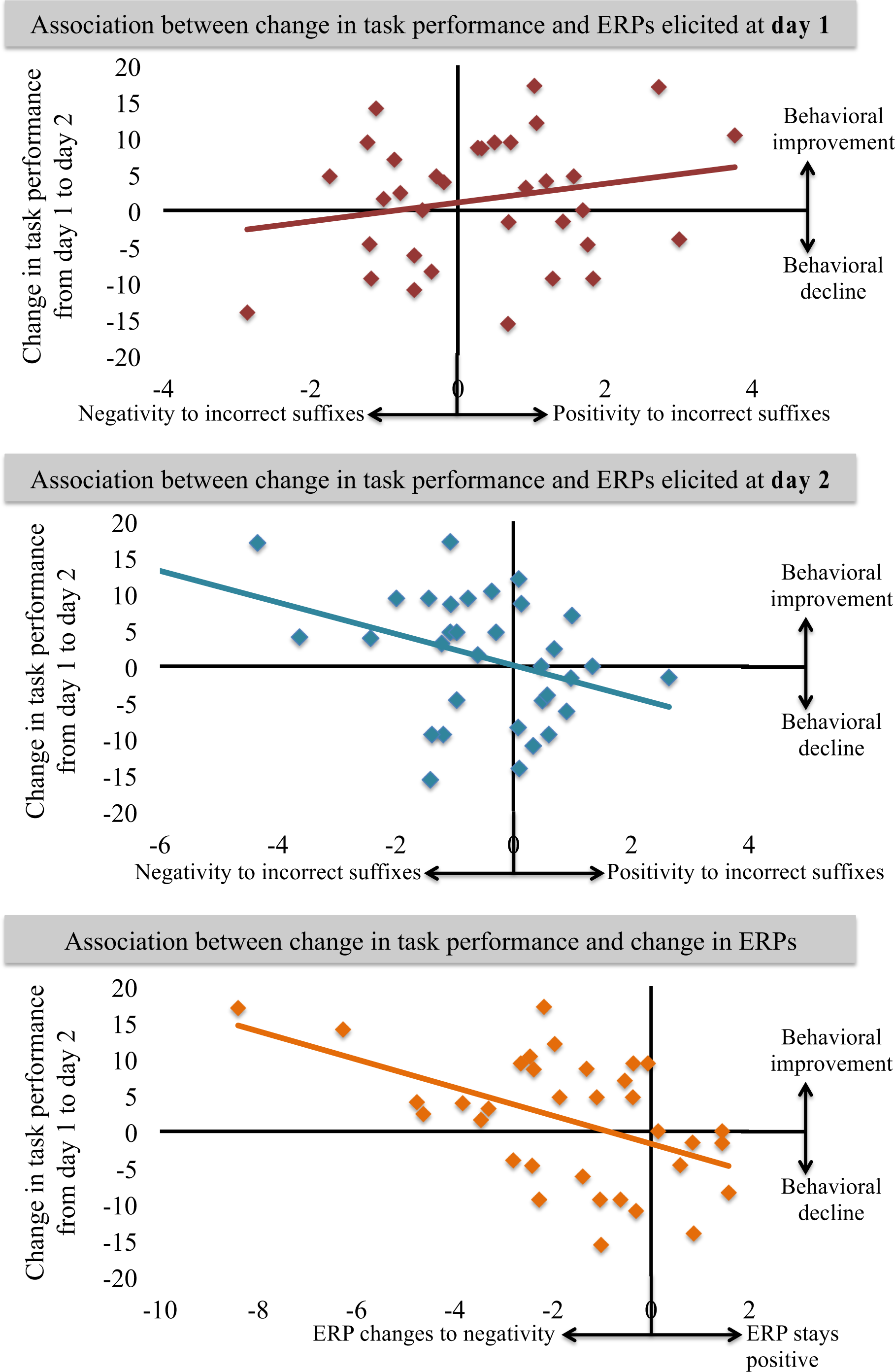
Association between change in task performance and event-related potentials (ERPs). The upper panel of the figure illustrates how the ERP in response to incorrect suffixes (i.e., ERP incorrect suffix – ERP correct suffix) on day one is related to the individual change in task performance (i.e., correctly answered trials in percent) from day one to day two (*r* =.28). The middle panel illustrates how the ERP to incorrect suffixes (i.e., ERP incorrect suffix – ERP correct suffix) on day two is related to the individual change in task performance (i.e., correctly answered trials in percent) from day one to day two (*r* = – .45). Children showing a more positive ERP to incorrect suffixes on day one are more likely to improve behaviorally compared to children showing a less positive ERP on day one. Similarly, children who showed a more negative ERP to incorrect suffixes on day two more strongly improved behaviorally from day one to day two compared to children who less strongly improved behaviorally. The lower panel illustrates how the change from a positive ERP on day one to a more negative ERP on day two is related to the individual change in task performance (*r* = –.52). Children showing a stronger change from positivity on day one to negativity on day two improved more strongly behaviorally from day one to day two.

Further, we calculated the difference between the positivity on day one and the negativity on day two, according to the procedure of calculating the change in task performance as described above. A negative value would indicate a stronger change from positivity on day one to negativity on day two. We again calculated Pearson’s bivariate correlation coefficient to now analyze the association between the changes in ERP polarity from day one to day two and the change in task performance from day one to day two.

The result showed a negative correlation between the individual change in ERP polarity from day one to day two and the individual change in task performance from day one to day two (*r* = – .52) (see Figure 4).

## 4. Discussion

The aim of the present ERP study was to investigate NAD learning by means of NAD violation recall using a miniature version of a natural language in 7-year-olds. Specifically, we not only tested NAD processing directly after learning, but also after a retention period involving sleep (i.e., at the next day). On the first day, German-speaking children were exposed to Italian sentences containing NADs (e.g. *La sorella sta cantando*; the sister is singing). Learning phases were followed by testing phases in which participants heard a mixture of correct sentences containing the same NADs as during learning phases, as well as incorrect sentences containing NAD violations (e.g. *La sorella sta cantare*; the sister is sing∅Φ; see Friederici et al., 2011; Mueller et al., 2009), while they performed a grammaticality judgment task. To then test recall of NADs, participants were re-invited the following day, on which they were presented with testing phases only, while again their EEG data and grammaticality judgments were acquired.

The grammaticality judgment task at either day did not reveal any NAD learning at the behavioral level (above-chance performance) in our group of 7-year-old children. This result was unexpected, given findings by Raviv and Arnon (2017), who showed that 7-year-olds had successfully learned an artificial grammar on a behavioral level. This discrepancy in results could have one of the following reasons: (1) Our natural language stimuli were more complex than the artificial language involving syllable triplets in Raviv and Arnon (2017)’s study and may thus be more difficult to learn. (2) Our grammaticality judgment task was more difficult than the two-alternative forced choice (2-AFC) task used by Raviv and Arnon (2017), in which children were presented with two stimuli, one of which conformed to a familiarized language and one did not. It is conceivable that having the direct comparison between a correct and an incorrect example in the 2-AFC task, including the knowledge that one sentence is correct and one incorrect, facilitates learning compared to the grammaticality judgment task used in the present study. Thus, it might be concluded that the present grammaticality judgment task is still too difficult for 7-year-old children (see also Lammertink, Witteloostuijn, Boersma, Wijnen, & Rispens, 2018) such that they cannot successfully show the same behavior as adults (i.e., above chance level, see Mueller et al., 2009). Based on these results, it might be concluded that our group of 7-year-old children did not learn the NADs *explicitly*. However, when looking at behavioral changes in performance (i.e., correct responses) from day one to day two, we found behavioral changes in the positive direction for some children (i.e., more correct answers on day two compared to day one), possibly indicating NAD learning at least for some of the 7-year-olds after a retention period involving sleep.

In the following, we will first discuss the ERP findings of NAD processing at the group level, before elaborating on our descriptive analysis of the association between behavioral and neurophysiological responses at the individual level. At the neural level, we found a more positive ERP response to NAD violations (i.e., 800 to 1000 ms after suffix onset) during testing phases on day one. In contrast, we found a more negative ERP response to NAD violations (i.e., 600 to 1000 ms after suffix onset) during testing phases on day two.

Regarding the latencies of the observed ERP components, the positivity on day one occurred slightly later than the negativity on day two. Since shorter latencies are typically interpreted as reflecting faster, more automatic processing (e.g. Friederici, Kotz, Werheid, Hein, & von Cramon, 2003), it is possible that the detection of NAD violations was still somewhat slower on day one and became faster and possibly more automatic after children had time to consolidate the learned NADs during sleep. Regarding the change in polarity of the ERP effects following overnight-retention, we suggest a change in representation as interpretation. In previous studies using this paradigm, positive ERPs in response to NAD violations have been reported for infants (Friederici et al., 2011) and adults when their PFC was inhibited by tDCS (Friederici et al., 2013). It has been suggested that infants employ associative learning strategies and that with increased PFC development, learning mechanisms change to controlled top-down learning (Skeide & Friederici, 2016). Because both, infants, whose PFC is not yet fully developed (Huttenlocher, 1990), and adults with a temporarily inhibited PFC, showed a positive ERP response to NAD violations, more positive ERPs have been interpreted to indicate associative learning of NADs (Friederici et al, 2011; 2013). In our study, the positivity on day one may thus indicate that children have formed associative representations of the NADs before retention. In contrast, we found a negative ERP to NAD violations on day two. Negative ERP responses to NAD violations in this paradigm have been reported for adults under standard conditions, that is, when their PFC was not inhibited (Friederici et al., 2013; Mueller et al., 2009; Citron et al., 2011). This negativity (occurring approx. 340 to 540 ms after suffix onset with a centro-parietal distribution) was interpreted to reflect an N400, indicating lexical access, based on perceptual features (Mueller et al., 2009; Kutas & Federmeier, 2000). Children’s negativity found in the present study on day two occurred slightly later (i.e., 600 to 1000 ms after suffix onset) and more frontally, most likely reflecting an immature N400 (see Hahne, Eckstein, & Friederici, 2004; Henderson, Baseler, Clarke, Watson, & Snowling, 2011), indicating a lexical strategy (Mueller et al., 2009) on testing day two. We speculate that children’s NAD representation may have initially been a phonological association stored in episodic memory. These associative representations were then most likely transferred to long-term memory overnight (comparable to how new words are learned and consolidated over night; see Davis & Gaskell, 2009; Henderson et al., 2012; Tamminen et al., 2010), which children then tried to access during testing phases on day two. Consequently, NAD violations would then have led to a larger negative ERP component due to retrieval difficulties. It is possible that these long-term representations of NADs were lexicalized, as in adults (Mueller et al., 2009), perhaps as whole phrases. Under this view, children would have attempted to store the dependent elements of the NADs as lexicalized phrases in lexical long-term memory, as indicated by an immature N400-like response. While it is likely a more efficient way to learn and store the NADs in an associative way, like infants do (Friederici et al., 2011), a lexical strategy has been proposed as the mechanism adults use to learn NADs (Mueller et al., 2009). Based on the similarity of ERP components in our study compared to adults’ ERPs (see Mueller et al., 2009), we propose that 7-year-olds employed this strategy on the second day. Further, it is likely that children employed an implicit rather than an explicit strategy, as we found significant ERP effects on both testing days, but no significant behavioral effects. Taken together, we suggest the positivity in the ERP on day one to indicate associative NAD learning and the negativity in the ERP on day two to indicate a lexical processing strategy, where both mechanisms might be beneficial for NAD learning in 7-year-old children.

ERPs have previously been used to study children’s NAD learning during infancy (Friederici et al., 2011) and early childhood (Mueller et al., 2018) and have been shown to have a strong test-retest reliability (Cassidy et al., 2012). In contrast, behavioral measures of individual differences in statistical learning, such as grammaticality judgments and 2-AFC tasks, have recently been criticized for being unreliable at the individual level (Siegelman et al., 2017a; West et al., 2018). Specifically, when tests that were developed for group-level inferences are used to study individual differences, they often do not have enough statistical power at the individual level. This can be due to several factors, such as 1) a low number of trials, 2) all trials having the same difficulty, and 3) group-level performance often being at chance-level (Siegelman et al., 2017a). In light of these concerns, we will discuss individual differences in ERPs in relation to mean change in behavior in the grammaticality judgment task from day one to day two in terms of a descriptive analysis in the following.

Both, the positivity on day one, as well as the negativity on day two, were associated with children’s behavioral changes in performance from day one to day two. Specifically, children who showed a more positive ERP response to NAD violations on day one and children who showed a more negative ERP response to NAD violations on day two showed a stronger behavioral change towards more correct answers on day two compared to day one. These results possibly indicate that 7-year-olds show NAD recall after a retention period involving sleep and that their representations of NADs change over this retention period (the ERP polarity change might indicate a representation change). Further, we showed that an individual’s stronger change from a positive ERP effect on day one to a negative ERP effect on day two was correlated with stronger positive behavioral changes (i.e., more correct answers on day two compared to day one). These results have to be interpreted with caution because individual differences in statistical learning show questionable reliability (Arnon, 2019; Siegelman et al., 2017a, 2017b; West et al., 2018) and the behavioral performance was not above chance at the group level. However, our results at the individual level are in line with our results at the group level. Both analyses lend support to the interpretation of a change of NAD representation during a retention period enabling 7-year-olds’ recall of NADs on the second testing day.

A study by Friedrich and colleagues (2017) offers insight into a possible mechanism underlying this change of representations during a retention period. Specifically, this study demonstrates an effect of sleep on the representation of learned associations between object-word pairs in infants. Object-word pairs were learned through mere phonological associations by infants who had a short nap after familiarization, while infants who had a longer nap built up semantic long-term memory representations of the object-word pairs. Infants who did not sleep between familiarization and test, however, did not show any evidence for learning the object-word pairs. Similarly, it is possible that in our study some children built up an associative representation of NADs on day one (indexed by a stronger positivity), possibly in episodic memory. The retention period between day one and day two then may have allowed those children to consolidate their associations and transfer them to long-term memory (in line with system consolidations theory, see e.g. Davis & Gaskell, 2009). This consolidation may have enabled children to build more robust representations (indexed by a stronger negativity) of the NADs, enabling these children to recall the NADs on day two, as indicated by more correct answers on day two. Our results are in line with previous studies reporting a beneficial effect of a consolidation period involving sleep for artificial grammar learning in infants (Gómez et al., 2006; Hupbach et al., 2009) and for word learning in older children (Backhaus et al., 2008; Henderson et al., 2012; Smith et al., 2016). More specifically, the idea that successful performance in an artificial grammar-learning task is only achieved after consolidation involving sleep is in line with a study by Fischer and colleagues (2006).

Crucially, only after a consolidation period that involved sleep, but not a consolidation period without sleep, did participants perform significantly above chance in the artificial grammar-learning task (Fischer et al., 2006). Our study provides further support for the beneficial effect of sleep on NAD learning.

It has recently been debated whether (artificial) grammar learning is governed by similarity-based or rule-based learning mechanisms (see Hahn & Chater, 1998). Similarity-based learning occurs when (chunks of) familiarized items are memorized and these memorized representations are then compared to test items during testing phases. During rule-based learning, on the other hand, abstract statistical regularities underlying the items are implicitly extracted and tested against the incoming test items. This would then result in stored rule-based mental representations. Opitz and Hoffmann (2015) provided evidence that both mechanisms play a role in artificial grammar learning, with similarity-based learning being especially prominent in initial stages of learning, while rule-based learning builds up over time. These different learning stages are supported by different brain structures with similarity-learning being sub-served by the hippocampus and rule-based learning by the PFC (Opitz & Friederici, 2003, 2004). In our study, it would be possible that on day one, children used similarity-based learning to associate the elements of the NADs on day one and that rule-based learning took place over the retention period allowing children to recall the NADs on day two. These different processing mechanisms would then account for the observed change from positivity to negativity in the ERPs. Because the change from positivity to negativity correlates with change in behavior from day one to day two, this might mean that only those children, who learned similarity-based on day one were able to transform their knowledge to rule-based representations on day two. However, in the present study, we cannot directly test whether children relied on similarity-based or rule-based representations at a given day and future studies will have to test this claim, for example by manipulating the instructions the participants receive. Specifically, with more explicit instructions regarding the underlying rules, participants could be nudged into employing rule-based rather than similarity-based mechanisms early on in learning.

### 4.1 Limitations

We examined the correlation of two measures of individual differences in NAD learning, one offline measure (change in behavior from day one to day two) and one online measure (ERPs). Since these measures were acquired in the same task, their correlation should not be over-interpreted. Recently, several concerns have been raised about the reliability of offline measures of individual differences in statistical learning (Siegelman et al., 2017a, 2017b; West et al., 2018; Arnon, 2019). In the present study, we address some of these concerns, as we used a relatively high number of trials (32 trials test phase trials per condition, which should allow for a relatively good discrimination between subjects’ NAD learning abilities; see Siegelman et al., 2017a). Moreover, we used ERPs, which have been shown to be strongly reliable (Cassidy et al., 2012). In addition, we found significant ERP effects at the group-level, supporting our interpretation of the identified individual differences (see also Siegelman et al., 2017a). However, we cannot account for all reliability issues raised in the literature. For example, there was no significant above-chance performance at the group level for the behavioral task, with most of the children performing at chance level on both days, making individual differences more difficult to interpret (see Siegelman et al., 2017a). The mean change in behavior from day one to day two was small, further making the interpretation of the mean change in behavior difficult. However, we believe that the association we found between ERPs and behavior supports the meaningfulness of change of behavior from day one to day two as a measure ^3^.

Another possible limitation of the present study is that we did not manipulate children’s sleep duration between day one and day two. Therefore, we cannot make claims about whether the change in NAD representations is specifically due to sleep or due to a more general effect occurring over the course of a consolidation period. Similarly, we did not manipulate whether or not children underwent testing phases on day one. Thus, we cannot clearly infer whether the differences between ERPs on day two (negativity) compared to day one (positivity) were due to repetition effects (i.e., testing phases on both days) or whether we would still find a negativity on day two if day one did not include a testing phase. Future studies will have to investigate whether the representational change also occurs in a sleep-wake design and when testing conditions are manipulated.

### 4.2 Conclusion

Taken together, the present study indicates that, even though 7-year-old children do not show above-chance level performance in our NAD grammaticality judgment task yet, NAD representations changed after a retention period including sleep, as indicated by ERP responses to NAD violations compared to familiarized NADs. At the group level, we found a more positive ERP response to NAD violations before retention, which could indicate associative representations of NADs and/or similarity-based learning. In contrast, after a retention period involving sleep, we found a more negative ERP response to NAD violations, indicating that representations had been stored in long-term memory and thus demonstrating recall of NADs. In a descriptive analysis of individual differences, a stronger change from a more positive ERP response on day one to a more negative ERP response on day two was associated with stronger positive changes in behavior (i.e., more correct answers on day two compared to day one). This could possibly indicate that only those children who had built associative representations of the dependencies on day one were able to consolidate these representations and show recall of NADs on day two, as indicated by a positive change in behavior (i.e., more correct answers on day two compared to day one). These results are the first to show children’s implicit recall of NADs embedded in a natural, foreign language after a retention period involving sleep.

## Acknowledgements

The Max Planck Society (GS, MP, RM, CM, ADF), the German Research Foundation, project FR 519/20-1 (FOR 2253; GS, MP, CM, ADF), and the Berlin School of Mind and Brain (MP) funded this project. The authors declare to have no conflict of interest. Further, we would like to thank all the participating families for their commitment, as well as Sophia Röthing for her help with setting-up the experiment and Ulrike Barth for her help with data acquisition and her dedicated work with the participating children.

## Footnotes

1 In order to do so, we used the formula 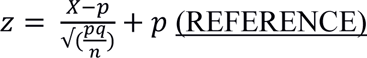. In this formula, refers to the child’s observed score, p refers to the probability of chance, q refers to the reciprocal of the probability of chance, and n to the total number of observations. We then set z to a critical value of ≥ 1.64 (i.e., the observed score is significantly different from chance at level p < .05, one-sided) and solved the equation to 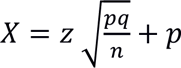 threshold indicating above chance-level performance.

2 Please note that other studies using the same stimuli (Citron et al., 2011; Friederici et al., 2013; Mueller et al., 2009) used visual inspection to identify relevant time windows, which we wanted to refrain from here, due to concerns of inflating the probability of type I errors.

3 Please note that our group-level analyses, on the other hand, are not affected by these concerns (see Arnon, 2019).

